# *Medicago truncatula* Zinc-Iron Permease6 provides zinc to rhizobia-infected nodule cells

**DOI:** 10.1101/102426

**Authors:** Isidro Abreu, Ángela Saéz, Rosario Castro-Rodríguez, Viviana Escudero, Benjamín Rodríguez-Haas, Marta Senovilla, Camille Larue, Daniel Grolimund, Manuel Tejada-Jiménez, Juan Imperial, Manuel González-Guerrero

## Abstract

Zinc is a micronutrient required for symbiotic nitrogen fixation. It has been proposed that in model legume *Medicago truncatula*, zinc is delivered in a similar fashion as iron, *i.e.* by the root vasculature into the nodule and released in the infection/differentiation zone. There, zinc transporters must introduce this element into rhizobia-infected cells to metallate the apoproteins that use zinc as a cofactor. *MtZIP6* (*Medtr4g083570*) is a *M. truncatula* Zinc-Iron Permease (ZIP) that is expressed only in roots and nodules, with the highest expression levels in the infection/differentiation zone. Immunolocalization studies indicate that it is located in the plasma membrane of rhizobia-infected cells in the nodule. Down-regulating *MtZIP6* expression levels with RNAi does not result in any strong phenotype when plants are being watered with mineral nitrogen. However, these silenced plants displayed severe growth defects when they depended on nitrogen fixed by their nodules, as a consequence of the loss of 80% of their nitrogenase activity. The reduction of this activity was not the result of iron not reaching the nodule, but an indirect effect of zinc being retained in the infection/differentiation zone and not reaching the cytosol of rhizobia-infected cells. These data are consistent with a model in which MtZIP6 would be responsible for zinc uptake by rhizobia-infected nodule cells in the infection/differentiation zone.

## INTRODUCTION

Plants require a steady supply of zinc (Frausto da Silva and Williams, 2001; Broadley et al., 2007). This element is an essential micronutrient used as cofactor by hundreds of enzymes in a typical cell, including representatives from the six major functional enzyme classes (Finkelstein, 2009). In addition, it also plays an important structural function in one of the largest families of transcription factors, the zinc finger proteins (Laity et al., 2001). Zinc, especially in alkaline soils, has low bioavailability leading to plant zinc deficiencies, and consequently, reduced yields and reduced nutritional value (Alloway, 2008; Wessells and Brown, 2012). As a result, a substantial research effort has been directed to discern how plants incorporate zinc from soil (Korshunova et al., 1999), how it is uploaded in the vasculature (Hussain et al., 2004), delivered to the phloem (Ishimaru et al., 2005; Yamaguchi et al., 2012), and carried to the embryo (Olsen et al., 2016). In addition, many of the soluble metal carriers and transcription factors regulating plant zinc nutrition have also been studied (Assunção et al., 2010; Sinclair and Krämer, 2012; Olsen and Palmgren, 2014). In contrast, little is known about how zinc reaches legume root nodules and its function therein. This is in spite of the importance that zinc has on symbiotic nitrogen fixation (SNF) (Ibrikci and Moraghan, 1993; O’Hara, 2001), one of the main pathways for nitrogen entry in the biosphere and a likely alternative to the overuse of polluting and expensive synthetic fertilizers (Herridge et al., 2008; Mus et al., 2016).

Legumes are able to establish an endosymbiotic relationship in their root nodules with a group of bacteria generally known as rhizobia (van Rhijn and Vanderleyden, 1995). These organs develop after a complex process triggered by bacterial and plant signals (Oldroyd, 2013; Downie, 2014). As a result, rhizospheric rhizobia penetrate into the root primordia following infection threads and are released into plant cells in an endocytic-like process (Catalano et al., 2006). Within these cells, rhizobia will differentiate into bacteroids surrounded by the plasmalemma-derived symbiosome membrane, and are then able to convert, fix, N_2_ into NH_4_^+^. In indeterminate type nodules, such as those presented by *Medicago, Vicia,* or *Pisum*, the developmental stages of the nodule can be followed along it, as different developmental zones (Vasse et al., 1990). This is due to indeterminate nodules having an apical meristem (Zone I) that grows as older parts are infected by rhizobia (early Zone II). In more mature regions of Zone II, rhizobia will differentiate into bacteroids by endoreduplications. In Zone III, the fixation zone, bacteroids will start fixing nitrogen, a process facilitated by the sudden loss of oxygen tension happening at the Interzone between Zones II and III (Soupène et al., 1995). Finally, in Zone IV, the nodule senesces and nutrients are recycled (Burton et al., 1998).

As expected from any endosymbiotic relationship, there is intense nutrient trafficking between the symbionts (Udvardi and Poole, 2013). The ammonia that has been produced in bacteroids is transferred to the host plant through the symbiosome membrane in exchange for photosynthates and mineral nutrients (phosphate, sulfur, iron, zinc…) that are provided by the host plant. In the case of iron, it has been shown that it is delivered by the vasculature and released in the apoplast of zone II in *M.truncatula* nodules (Rodríguez-Haas et al., 2013), where a plasma membrane transporter introduces the metal into rhizobia-infected cells (Tejada-Jiménez et al., 2015). A similar mechanism has been proposed to occur for other transition micronutrients, zinc among them (Rodríguez-Haas et al., 2013; González-Guerrero et al., 2014). If this is true, a zinc transporter must exist in the plasma membrane of Zone II cells, that when mutated affects zinc uptake by rhizobia infected cells.

Candidate transporters to carry out apoplastic zinc uptake would belong to the Yellow Stripe-Like (YSL), or to the Zinc-Iron Permease (ZIP) families. Phenotypical and biochemical studies indicate that proteins from these groups introduce divalent transition metals (Zn^2+^ among them) into the cytosol of the cell (Guerinot, 2000; Curie et al., 2008). YSLs do this as metal-nicotianamine complexes (DiDonato et al., 2004), while no type of complex seems to be necessary for transport by ZIP proteins (Lin et al., 2010). Given that iron uptake by rhizobia-infected cells is mediated by a Nramp family member (Tejada-Jiménez et al., 2015), transporters that do not use metal-nicotianamine as substrate (Nevo and Nelson, 2006), by analogy, it could be hypothesized that the more likely candidate for zinc uptake would be a ZIP transporter. This is further supported by transcriptomic data that shows that the ZIP transporter *MtZIP6* (*Medtr4g083570*), is highly expressed in rhizobia-infected cells (Limpens et al., 2013). In addition, another ZIP protein has already been associated with SNF, as being responsible for maintaining zinc homeostasis in the symbiosome in *Glycine max* (Moreau et al., 2002; Clarke et al., 2014).

In this study, *MtZIP6* was further characterized. The data support that it is responsible for zinc uptake by rhizobia-infected cells in *M. truncatula* nodules. *MtZIP6* expression peaks in the apical parts of the nodule, where its encoded protein is inserted in the plasma membrane. Although it was reported to be a Zn^2+^ and Fe^2+^ uptake transporter in yeast (López-Millán et al., 2004), *in planta* its role seems to be confined to zinc homeostasis in the context of SNF, since no phenotype was observed under non-symbiotic conditions, and only zinc homeostasis was affected in nodulated *mtzip6* RNAi plants. This work adds to our understanding of transition metal nutrition and homeostasis in legume nodule cells.

## RESULTS

### *MtZIP6* is highly expressed in nodules

Real time quantitative PCRs were used to determine *MtZIP6* expression in roots and shoots from non-inoculated, nitrogen-fertilized *M. truncatula* R108 plants, and in roots, nodules, and shoots from plants inoculated with *Sinorhizobium meliloti* (Fig. 1A). No transcripts were detected in shoots either in inoculated or non-inoculated plants. In roots, nodulation appeared to cause no significant change in transcript levels. However, the expression maximum was reached in nodules, with over four times the levels detected in nitrogen-fertilized, non-inoculated roots. Sequence comparison of the 16 ZIP proteins encoded in *M. truncatula* genome with those from other plants show that there is little correlation between sequence similarity and substrate preference for a determined divalent transition metal within the ZIP family (Fig. 1B). Contrary to what was expected, MtZIP6 shares little similarity to GmZIP1, a ZIP family member reported to be nodule-specific in *G. max* (Moreau et al., 2002). This suggests that both transporters could play different roles in the nodules. This is further indicated by their different substrate preference in yeast, since GmZIP1 transports Zn^2+^ towards the cytosol, while MtZIP6 also transports Fe^2+^ in the same direction (López-Millán et al., 2004). Figure 1B also shows that MtZIP6 shares little relationship with the other fifteen ZIP family members in *M. truncatula*, being MtZIP16 the closest one (72 % similarity).

**Figure 1.**
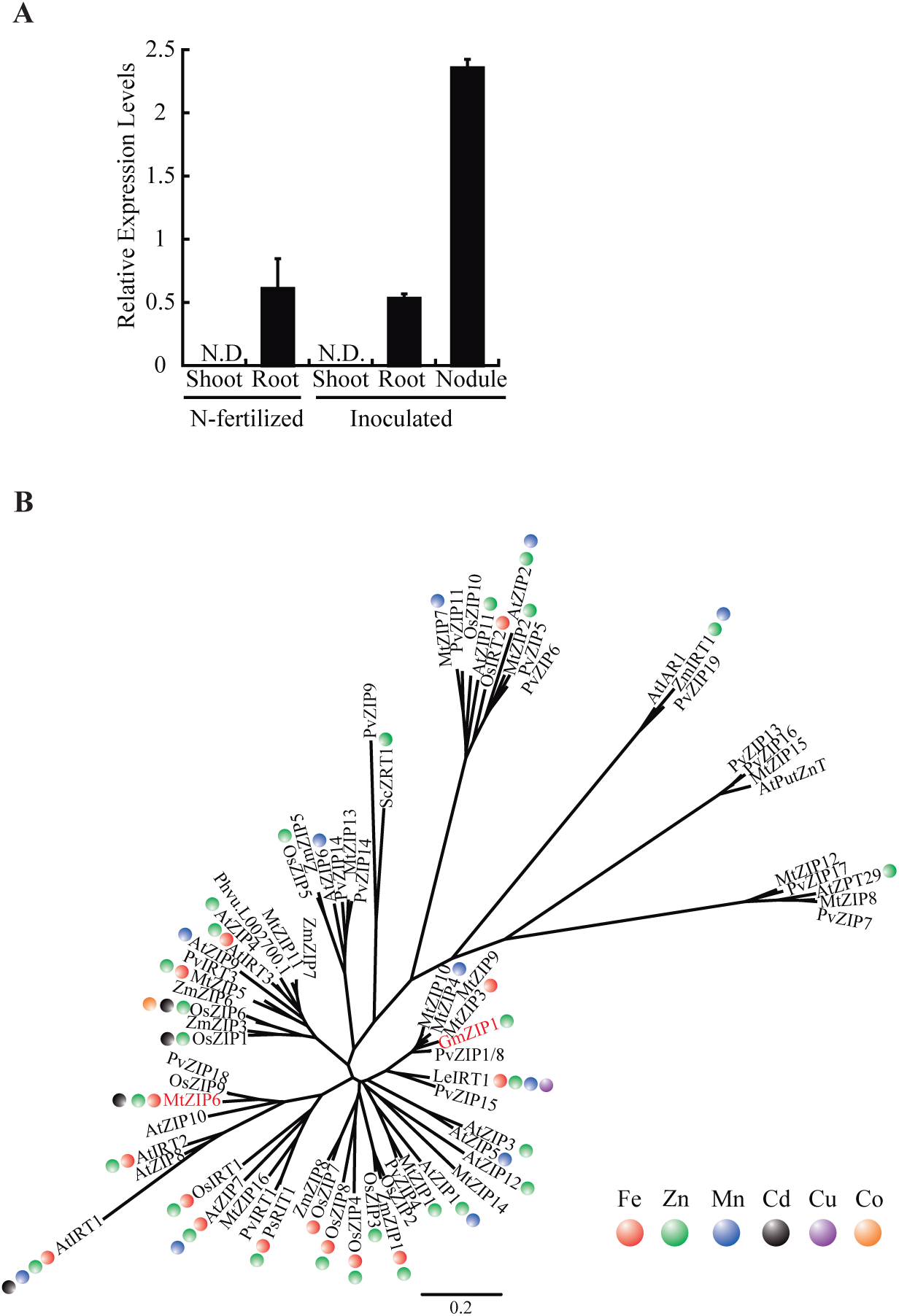
*MtZIP6* gene expression and relationship to other plant ZIP family members. A, *MtZIP6* expression in roots, shoots and nodules relative to internal standard gene *ubiquitin carboxyl-terminal hydrolase*. Data are the mean ± SD of three independent experiments. N.D. indicates that transcripts were undetectable. B, Unrooted tree of *M. truncatula* ZIP transporters, and representative plant ZIP homologues (see Material and Methods). Colored circles indicate proposed metal substrate according to literature.

### MtZIP6 is localized in the plasma membrane of cells in the infection/differentiation zone of the nodule

Metal release from the vasculature into the nodule appears to be primarily carried out in the differentiation zone of the nodule (Rodríguez-Haas et al., 2013). There, transporters must translocate the metals into the rhizobia-infected cells. Should MtZIP6 be carrying out this function, it will primarily be expressed in the proximities of the differentiation zone and the following cell layers. To test this, β-glucuronidase(GUS) activity was determined in *M. truncatula* plants expressing the *gus* gene under the transcriptional control of a DNA fragment containing 2 kb upstream from *MtZIP6* start codon. The reporter assays indicate that *MtZIP6* is primarily expressed in the apical regions of the nodule and in the root vasculature (Fig. 2A). Sections of these organs showed that in nodules the expression was located in the differentiation zone (Zone II) and in the younger parts of the fixation zone (Zone III) (Fig 2B). This expression pattern along the different nodule zones is in agreement with the transcriptomic data from the Symbimics database (Roux et al., 2014). These data were obtained from RNA extracted from laser-capture microdissected *M. truncatula* nodule cells from the Zone I, early and late Zone II, interzone II-III, and Zone III. Figure 2C shows that the majority of the transcripts are detected in the older part of Zone II, the interzone and Zone III, the areas with the maximum GUS activity (Fig. 2C). In the roots, *MtZIP6* is expressed in the vasculature around the xylem (Fig 2D).

**Figure 2.**
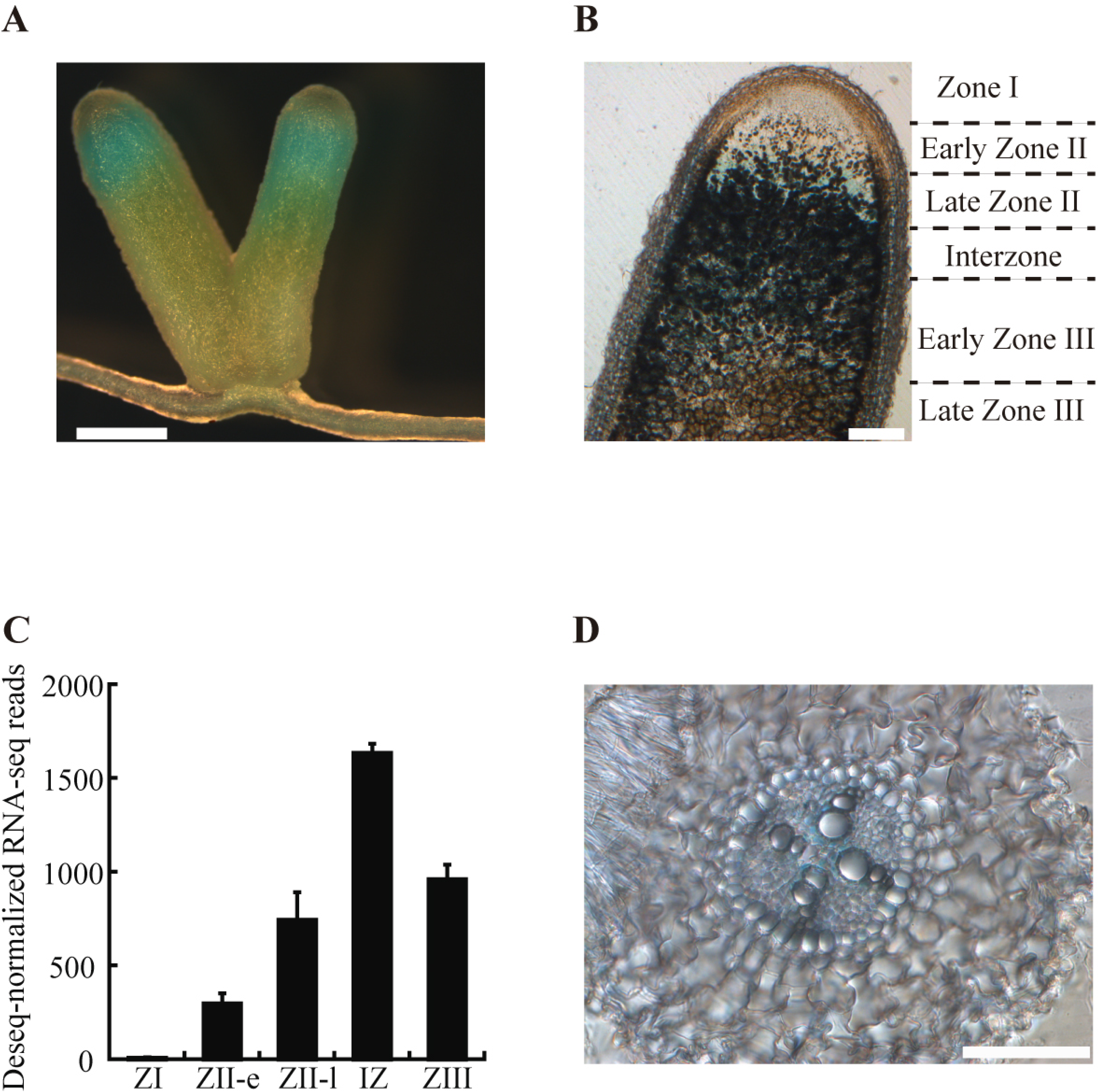
Tissular localization of *MtZIP6* expression. A, Histochemical staining of GUS activity in the root and nodules of *M. truncatula* plants transformed with plasmid pGWB3 containing the *MtZIP6-*promoter:*gus* fusion. Scale bar represents 1 mm. B, Longitudinal section of GUS stained nodule from *M. truncatula* plants transformed with plasmid pGWB3 containing the *MtZIP6-*promoter: *gus* fusion. Scale bar represents 200 μm. C, Expression of *MtZIP6* in the different nodule zones: ZI indicates Zone I; ZII-e, early Zone II (closest to Zone I); ZII-l, late Zone II; IZ is the interzone between Zones II and III; and ZIII, Zone III. Data were obtained from the Symbimics database (https://iant.toulouse.inra.fr/symbimics). D, Cross section of a GUS-stained root from from *M. truncatula* plants transformed with plasmid pGWB3 containing the *MtZIP6-*promoter::*gus* fusion. Scale bar represents 100 μm.

Further validation of these results was obtained in plants transformed with *MtZIP6,* whole gene structure (introns+exons) under the control of the same promoter region used in the GUS expression assays, fused to three hemagglutinin (HA) epitopes in C-terminus (MtZIP6-HA). Addition of this tag had no major effect on the functionality of the protein, since both HA-tagged and un-tagged MtZIP6 could complement the zinc uptake phenotype of yeast mutant *zrt1/zrt2* (Suppl. Fig. S1). MtZIP6-HA was detected using an Alexa594-conjugated antibody, and its position within the nodule was mapped with the help of DAPI, to stain DNA, and a constitutive GFP-expressing *Sinorhizobium meliloti*. As expected, the immunolocalization studies supported the histochemistry results, showing that MtZIP6-HA was primarily located in the differentiation zone of the nodules (distal ZII - apical ZIII). Alexa594 signal was not detected in older parts of the fixation zone, nor in the meristematic or in the infection areas (Fig. 3A). At the cellular level, MtZIP6 was detected in the periphery of the rhizobia-infected cells, indicative of the plasma membrane or a region close to it (Fig. 3B). No MtZIP6 was detected in rhizobia-free cells in Zone II. To confirm the putative plasma membrane localization, *MtZIP6* coding sequence was fused to a green fluorescent protein (GFP) tag under the control of a 35S promoter and co-agroinfiltrated into *Nicotiana benthamiana* leaves with a cyan fluorescent protein (CFP)-labelled plasma membrane marker. Those cells that expressed both constructions showed an overlap of the GFP and CFP signals, consistent with a colocalization in the plasma membrane (Suppl. Fig. S2). In roots, as already indicated by the GUS expression assays, MtZIP6-HA was localized in the stele, in close proximity to the xylem (Fig. 3C).

**Figure 3.**
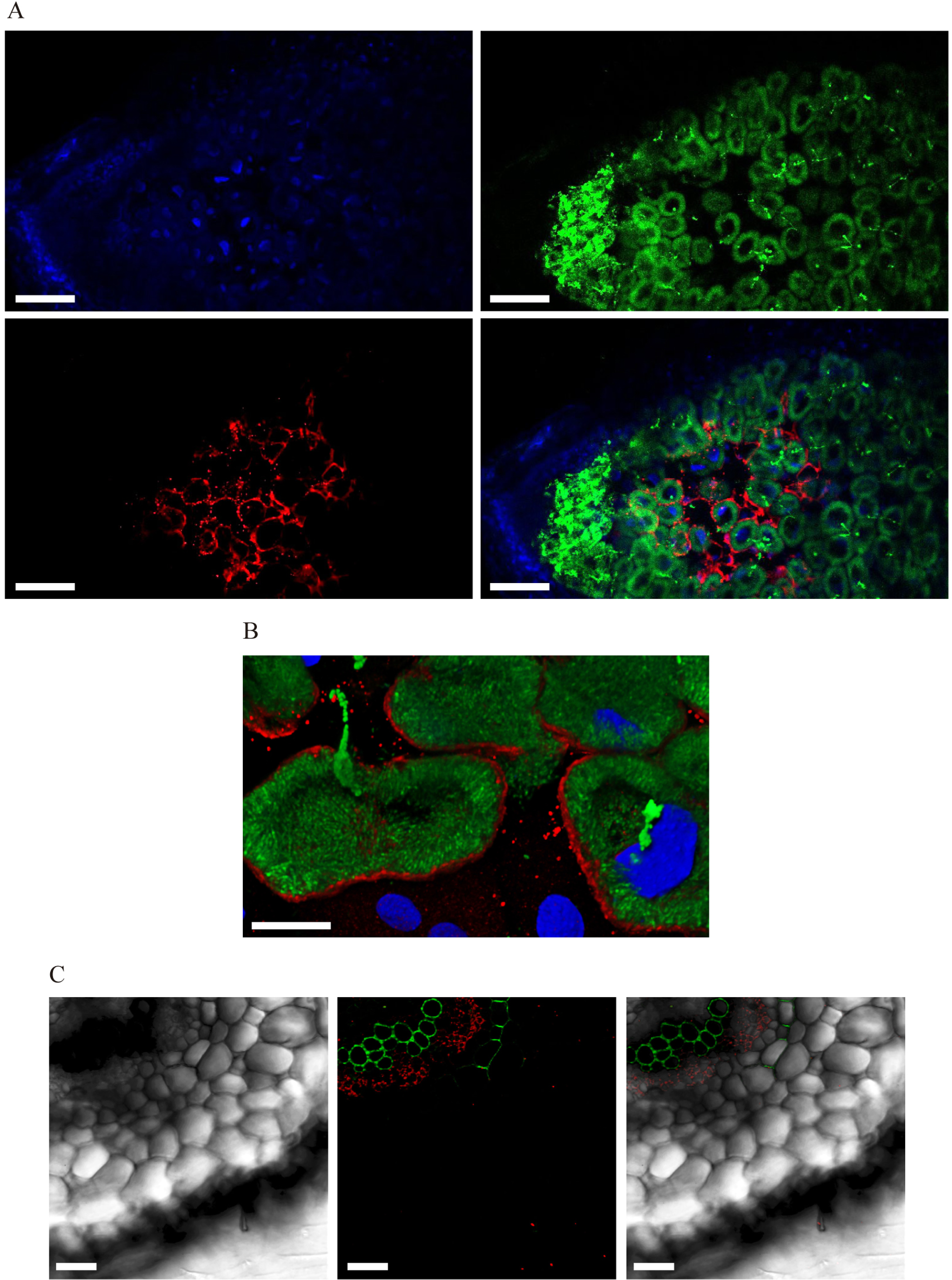
Subcellular localization of MtZIP6-HA. A, Cross section of a 28-dpi *M. truncatula* nodule inoculated with *S. meliloti* constitutively expressing GFP (green, upper right panel) and transformed with a vector expressing the fusion MtZIP6-HA under the regulation of its endogenous promoter. Nodules were stained with DAPI to show DNA (blue, upper left panel). MtZIP6-HA localization was determined using an Alexa 594-conjugated antibody (red, lower left panel). The lower right panel shows the overlay of DNA, *S. meliloti*, and MtZIP6-HA. Scale bar represents 100 μm. B, Three-dimensional reconstruction of MtZIP6-HA expressing cells. GFP-expressing *S. meliloti* are shown in green, red indicates the position of MtZIP6-HA, and blue is DAPI-stained DNA. Scale bar represents 25 μm. C, Cross section of a *M. truncatula* root transformed with a vector expressing the fusion MtZIP6-HA under the regulation of its endogenous promoter. Left panel shows the transillumination figure. Middle panel shows MtZIP6-HA localization using an Alexa 594-conjugated antibody (red) and xylem and endodermis layers using autofluorescence (both green). Right panel shows the overlaid images. Scale bar represents 50 μm.

### *MtZIP6* silencing has a negative effect on symbiotic nitrogen fixation

To study the role of *MtZIP6* in plant metal nutrition and in SNF, RNAi plants were produced by *A. rhizogenes* transformation. Given that *MtZIP6* was not expressed in shoots, hairy-root transformed plants would be sufficient to study the effect that down-regulating *MtZIP6* has on the plant. However, as no germ-line transformants were produced, and given that gene silencing can vary from one line to another, pooled plants were used in these analyses to average the effects of each independent T-DNA insertion event. Using the first 469 bp of *MtZIP6* cDNA to silence the gene resulted in an average 80% reduction of expression (Fig. 4A). No reduction on transcript levels was observed in any of the three ZIP family members with the highest degree of identity to *MtZIP6* (*MtZIP5, MtZIP11,* and *MtZIP16*) (Suppl. Fig. S3) Under non-symbiotic conditions, when plants were not inoculated and the nutritive solution was supplemented with NH_4_NO_3_ no alteration in growth was observed between *mtzip6* RNAi plants and controls which express 450 bp of the *gus* gene in the same vector as the RNAi plants (Fig. 4B). Similarly, no significant differences were observed when comparing shoot and root dry biomass between control and RNAi plants (Fig. 4C).

**Figure 4.**
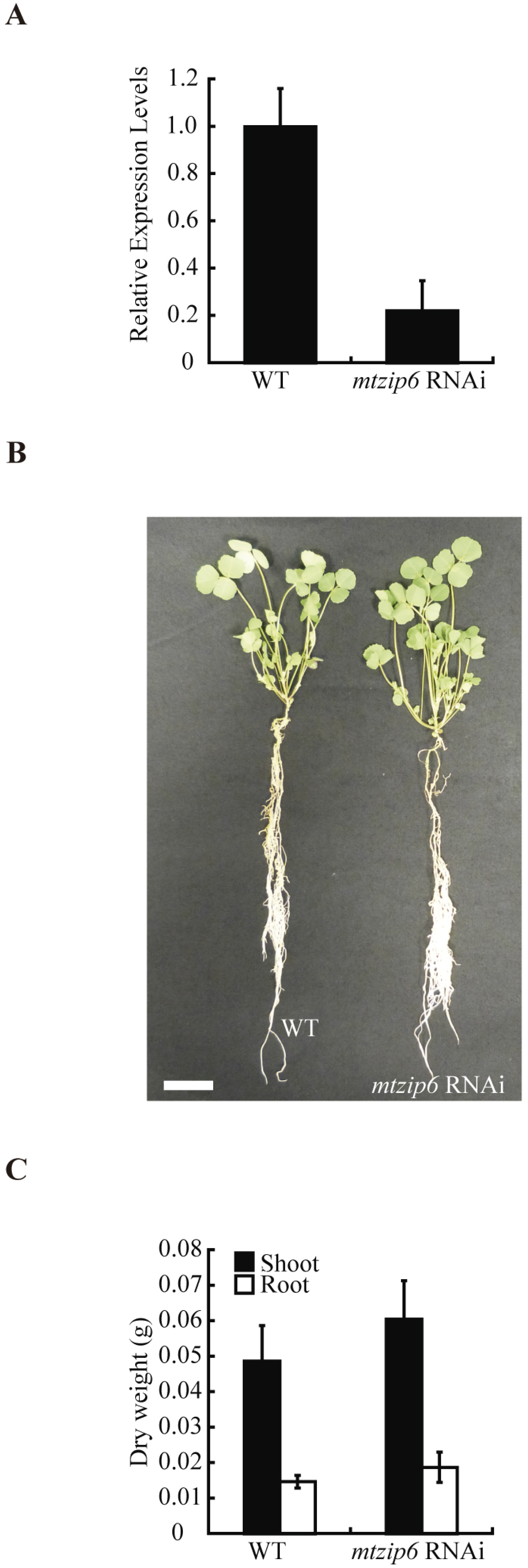
Phenotype of *mtzip6* RNAi plants under mineral N supply conditions (non-symbiotic). A, Expression levels of *MtZIP6* in 28 dpi nodules from control (WT) and *mtzip6* RNAi plants relative to internal standard gene *ubiquitin carboxyl-terminalhydrolase* and relative to control expression levels. Data are the mean ± SD of three independent experiments. B, Growth of representative wild type and *mtzip6* RNAi plants. Scale bar represents 3 cm. C, Dry weight of shoots and roots. Data are the mean ± SD (n= 10 plants).

In contrast, when the plants were inoculated with *S. meliloti* and no fixed nitrogen was provided in the nutritive solution used to water them, plant growth was severely affected in *mtzip6* RNAi plants (Fig. 5A). These plants showed reduced biomass production, with an average reduction of 70% in shoots and 68% in roots (Fig. 5B). Nodule development was affected, with approximately half of the nodules smaller and, white, likely non-functional (Fig. 5C-D). These observations are the likely result of an impairment in nodule function, with a subsequent reduction in nitrogenase activity. To measure this activity, the acetylene reduction assay (Hardy et al., 1968) was used. On average, RNAi plants showed a 80% reduction in nitrogenase activity per plant when compared to control ones (Fig. 5E), which was not due to a reduction in the nitrogenase protein (Suppl. Fig S4). These results obtained from pooling plants were consistent with the average of analyzing individual plants resulting each from one independent transformation event (Suppl. Fig S5)

**Figure 5.**
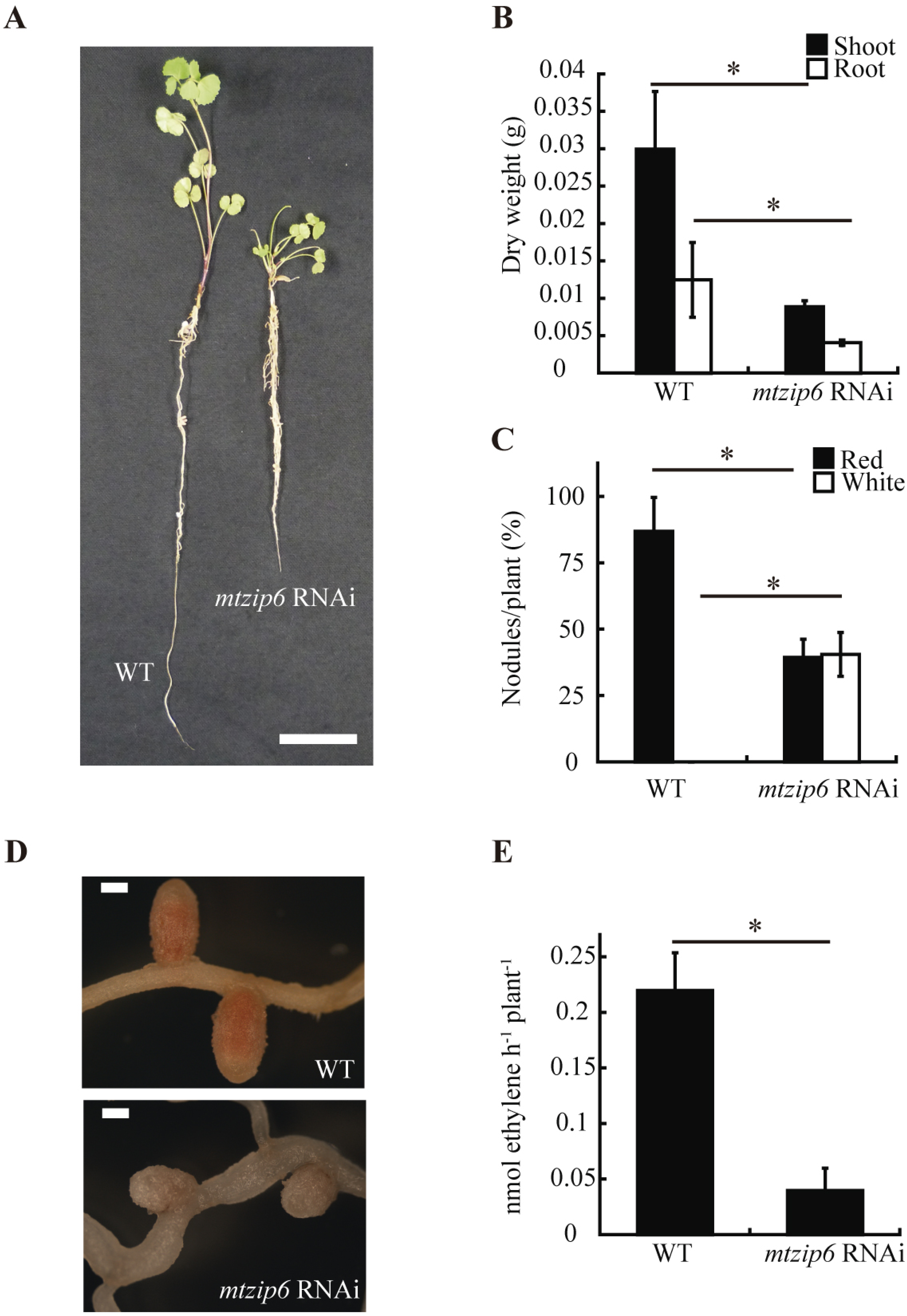
Symbiotic phenotype of *mtzip6* RNAi plants. A, Growth of representative wild type and *mtzip6* RNAi plants. Scale bar represents 3 cm. B, Dry weight of shoots and roots. Data are the mean ± SD (n= 11-12 plants). C, Nodule number per plant. 100% = 3.48 nodules/plant. Data are the mean ± SD (n= 20 plants). D, Close view of a representative wild type nodule and a white nodule. Scale bar represents 500 mm. E, Nitrogenase activity in 28 dpi nodules. Data are the mean ± SD of three sets of five pooled independently transformed plants. Asterisk indicates significant differences (P ≤ 0.05).

### MtZIP6 provides zinc to rhizobia-infected nodule cells

To determine the substrate of MtZIP6, metal content in shoots, roots, and nodules in wild-type and *mtzip6* RNAi plants was determined. While no change was observed for iron levels (Suppl. Fig. S6), there was an increase on zinc content in nodules of *mtzip6* RNAi plants (Fig. 6A). This result was indicative of a possible role of MtZIP6 in zinc homeostasis, which was also supported by the zinc-dependent regulation of *MtZIP6* expression in roots (Fig. 6B). *MtZIP6* transcription in nodules was not affected by zinc concentrations in the nutritive solution. However, in spite of low zinc-levels upregulating *MtZIP6*, no difference in biomass production or nitrogenase activity were observed between wild-type and RNAi plants when grown under low-zinc conditions (Suppl. Fig. S7). Similarly, increasing zinc concentrations in the nutritive solution by 10 or 100-fold did not result in a restoration of a wild type phenotype in *mtzip6* RNAi plants (Suppl. Fig. S8), and increasing them by 500-fold was equally toxic for wild-type and silenced plants (data not shown).

**Figure 6.**
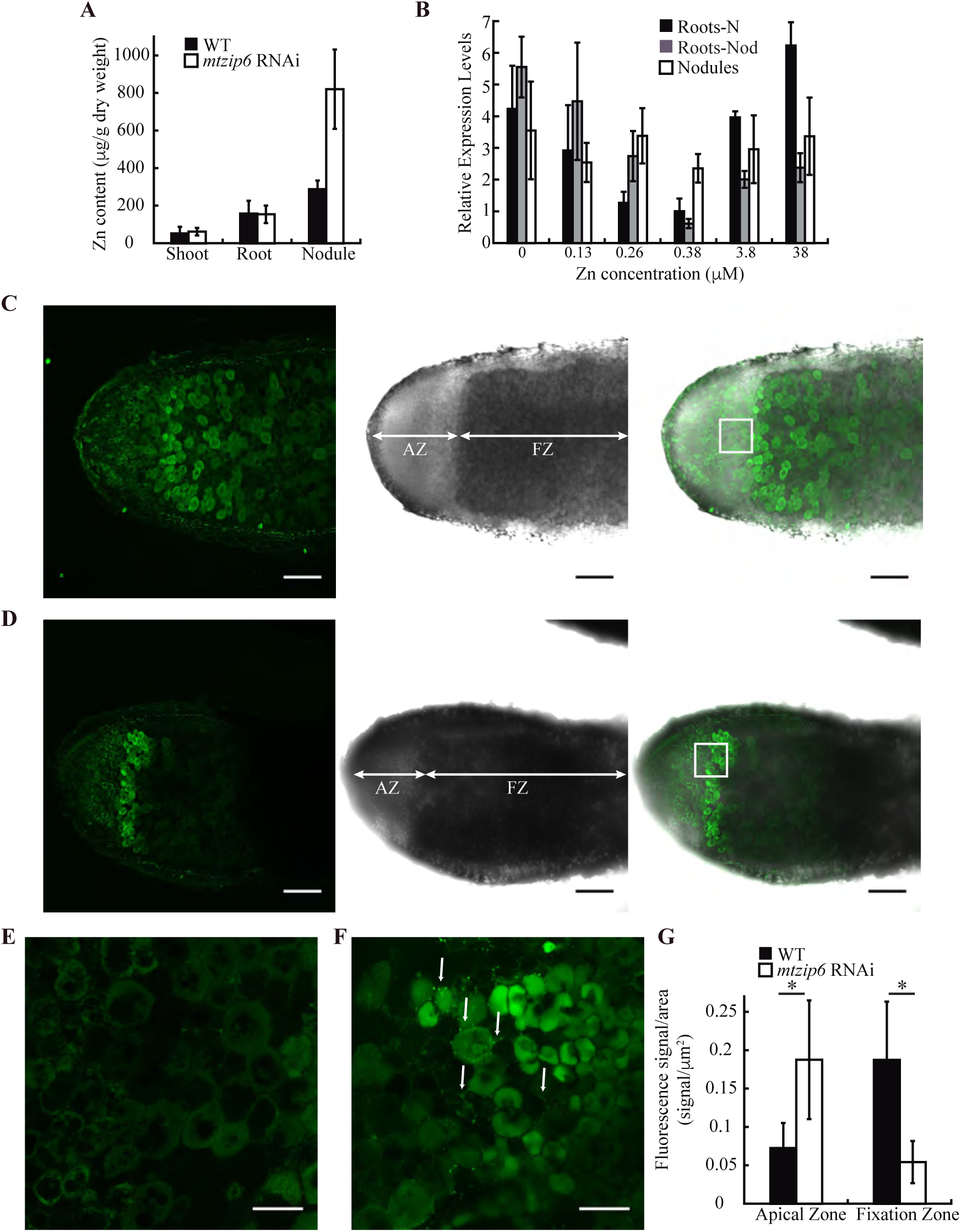
Role of MtZIP6 in zinc homeostasis. A, Zinc content (ppm) in 28-dpi plants. Data are the mean ± SD of two sets of five pooled transformed plants. B, Effect of zinc concentration on *MtZIP6* expression in roots from nitrogen-fertilized plants and roots and nodules from inoculated ones relative to internal standard gene *ubiquitin carboxyl-terminalhydrolase*. Data are the mean ± SD of three independent experiments. C, Zinc distribution in control nodules determined with Zinpyr-1 (green, left panel), transillumination (middle panel) and overlay (right panel). AZ and FZ indicate apical and fixation zones, respectively. D, Zinc distribution in *mtzip6* RNAi nodules determined with Zinpyr-1 (green, left panel), transillumination (middle panel) and overlay (right panel). AZ and FZ indicate apical and fixation zones, respectively. E, Detail of zinc distribution in the apical zone of control nodules. F, Detail of zinc distribution in the apical zone of *mtzip6* RNAi nodules. Arrows indicate apoplastic zinc deposits. G, Quantification of Zinpyr-1 signal per area in the apical and fixation zones of control and *mtzip6* RNAi nodules. Data are the mean ± SD (n=5-12 nodules). Scale bar in C-D represents 200 μm. Scale bar in E-F represents 50 μm. Asterisk indicates significant differences (P ≤ 0.05).

To determine whether silencing *MtZIP6* has an effect on zinc distribution, the localization of this element was determined using the zinc-sensitive probe Zinpyr-1 (Sinclair et al., 2007). The images analyzed showed that zinc content in wild-type nodules is higher in Zone III, in the same ring-like distribution that symbiosomes have in the cell (Fig. 6E). Silencing *MtZIP6* had no major effect in the distribution of zinc in the nodule (Fig. 6D), although a lower zinc content in Zone III was observed. At closer magnification, control plants showed very little zinc in the apoplast of late Zone II-Interzone (Fig. 6E). However, at closer magnification of the late Zone II-Interzone, several zinc deposits in the apoplast were detected in *mtzip6* RNAi nodules (Fig. 6F). This indicates that the increased zinc levels detected in *mtzip6* RNAi nodules could be the result of a zinc accumulation in the younger parts of the nodule, prior to the region where MtZIP6 is located. To test this, Zinpyr-1 signals were integrated before and after the interzone of several different nodules and standardized to surface units (selected images are shown in Suppl. Fig. S9). As it is shown in Figure 6G, the apical region of the nodule, corresponding to Zones I and II, had a significantly higher zinc content in RNAi plants than in their controls, while Zone III showed a significant reduction consistent with a reduction in the capacity to translocate zinc into rhizobia-infected cells. To ensure that these observations were not biased by the relative affinities of Zinpyr-1 towards zinc-binding proteins, zinc distribution was also assessed in freeze-dried nodule sections using synchrotron-based X-ray fluorescence (Suppl. Fig. S10).

To rule out a potential role of MtZIP6 as iron transporter in infected cells, we analyzed iron distribution and iron complementation in *mtzip6* RNAi plants. As with zinc, increasing iron concentrations in the nutritive solution did not restore the wild-type phenotype in *mtzip6* RNAi (Suppl. Fig. S6B). Iron distribution did not change in *mtzip6* RNAi plants compared to wild-type ones, and no significant differences were observed in the apoplast (Suppl. Fig. S6C and D). In addition, *MtZIP6* expression wasn’t altered in *nramp1-1*, a mutant altered in iron nodule homeostasis (Tejada-Jiménez et al., 2015) and complementary *MtNramp1* expression didn’t suffer any change in *mtzip6* RNAi plants (Suppl. Fig. S11).

## DISCUSSION

Zinc is a limiting nutrient for plants growing in many parts of the world (Alloway, 2008), with severe negative effects on growth, crop yields, and nutritional value (Wessells and Brown, 2012; Bashir et al., 2013). This is even more so in legumes, since they have additional sink organs, the nodules, that demand relatively large amounts of metallic micronutrients (O’Hara, 2001; Brear et al., 2013; González-Guerrero et al., 2014; González-Guerrero et al., 2016). Consequently, the nutritive properties and yields of legumes, one of the main vegetable protein sources in the world (Boye et al., 2010), greatly rely in our ability to better understand and improve zinc delivery to nitrogen fixation sites. This would also impact the design of future strategies based in ensuring the adequate supply of essential-limiting micronutrients to improve SNF capabilities in legumes or to introduce them in other crops. Using our knowledge on iron delivery to *M. truncatula* nodules as a model, we hypothesized that: i) Zinc is delivered by the vasculature and released in the apoplast of the differentiation zone of the nodule, and ii) a ZIP family member that is located in the plasma membrane of cells in this zone of the nodule is responsible for taking up this micronutrient. The results presented here support these hypotheses and point to MtZIP6 as the transporter carrying out this role.

Previous studies had shown that MtZIP6 is a functional ZIP transporter that is able to introduce iron and zinc into yeast cells (López-Millán et al., 2004). This protein was expressed in roots and in the apical region of the nodule. MtZIP6 was located in the plasma membrane of the rhizobia-infected cells in the differentiation zone of the nodule as indicated by transcriptomics, histochemistry and immunolocalization. Moreover, when *MtZIP6* transcript levels were down-regulated by RNAi, nitrogen fixation was significantly reduced. All these data support a role of MtZIP6 in SNF, but they do not indicate the nature of the substrate being transported *in planta*, whether it is introducing zinc into the cell or it is a functional analogue of MtNramp1 involved in iron metabolism (Tejada-Jiménez et al., 2015).

The available evidence indicates that *in planta* MtZIP6 works primarily as a zinc transporter. Its ability to introduce iron into yeast cells could simply be the result of overexpressing a transporter with a low iron affinity, that when pressed would transport iron, a phenomenon that is not unusual when assaying heterologous expression in yeast. Moreover, if MtZIP6 were involved in iron transport, we would expect to see an alteration of iron levels in the plant or nodules or a restoration of wild-type growth by watering with added iron, as it happens in *mtnramp1-1* or in *ljmate1* plants, both mutants affected in iron homeostasis in nodules (Takanashi et al., 2013; Tejada-Jiménez et al., 2015). Instead, an accumulation of zinc was detected in nodules of *mtzip6* RNAi plants, localized to the apical regions of the nodule, where apoplastic zinc deposits were observed. This would be consistent with a model in which reducing the levels of MtZIP6 would impede zinc transport across the plasma membrane, causing its accumulation in the apoplast and in the nodule region prior to where MtZIP6 is located. Our results also indicate that there must be some systemic signal indicating that zinc is limiting within the cell and that more of this element is required, which would explain why nodule zinc concentrations increase in the *mtzip6* RNAi nodules rather than remaining equal or even lower. Similar observations have been made when studying two *M. truncatula* mutants in nodule-specific molybdate and copper transporters (unpublished results). In addition, the expression pattern of *MtZIP6* and *MtNramp1* does not support the hypothesis that they have a complementary role in the uptake of the same metal. If they were carrying the same micronutrient, some type of co-regulation would be expected, in which when one is active, the other would be down-regulated, to prevent unnecessary protein translation. However, the opposite has been observed: reducing the expression levels of one of them (by mutation or by RNAi) results in down-regulation of the other. Consequently, the observed reduction in nitrogenase activity is not the direct result of a reduction of the delivery of a direct metal cofactor, but rather of the loss of one or several zinc-dependent functions that will affect nodule functioning. Pin-pointing the specific cause is extremely difficult with our current understanding of the nodule zinc-proteome. Since MtZIP6 is merely introducing zinc in the cytosol, a process that if diminished would affect the wide range of zinc-using proteins (enzymes, transcription factors, structural proteins), the observed phenotype could be the result of an overall compromise of the cell metabolism, affecting multiple cellular processes, including nitrogen fixation. Of all the possible causes, we think it is unlikely that a bacterial zinc-dependent transcription factor is involved, since nitrogenase protein levels are not altered in wild-type and RNAi plants. More work needs to be done in characterizing the nodule zinc-proteome to better ascertain the molecular basis of the phenotype observed.

*MtZIP6* is also expressed in the root vasculature, in a position consistent with the xylem parenchyma. However, the results obtained indicate that a reduction of 90% in *MtZIP6* expression is not sufficient to confer any altered phenotype when plants are being watered with an ammonium nitrate-supplemented nutritive solution. This could be the result of a stable mRNA that would allow for enough protein synthesis to fulfill its role or of some other transporter partially complementing MtZIP6 function in the root vasculature. In addition, the phenotype observed for *mtzip6* RNAi plants under symbiotic conditions is not the consequence of the loss of function of MtZIP6 in the root vasculature. If this were the case, we would detect less zinc reaching the nodules. However, the opposite is observed, with zinc localization studies showing that this metal is being delivered to the nodule apical region, but it is being retained there due to the reduction of the zinc uptake function of a knocked-down MtZIP6. This distribution pattern also supports a model in which other metals, and not just iron, are released by the vasculature in the apoplast of Zone II, and indicates that this region is critical for nodule metal nutrition.

It is also interesting to point out that MtZIP6 is only expressed in rhizobia-infected cells, a result supported by transcriptomic analyses of laser-captured microdissected cells (Limpens et al., 2013), raising the question of how zinc is taken up by non-infected cells. Being an essential micronutrient, these cells have to obtain zinc, and the likely candidates would be YSL or ZIP family members as indicated in the introduction. If it is a ZIP transporter, it will not be MtZIP6, and we would expect the other transporter to have a different affinity for zinc, so that it could be differentially distributed between the two cell types. Putative candidates for zinc uptake by non-infected cells are MtZIP1, MtZIP4 or MtZIP7, ZIP transporters that are mainly expressed in non-infected nodule cells (Limpens et al., 2013). Alternatively, if zinc uptake by non-infected cells is mediated by a YSL transporter, a prerequisite for it is the requirement of nicotianamine for nodule functioning, something that has been recently shown (Avenhaus et al., 2016).

In conclusion, as summarized in the model presented in Figure 7, zinc would reach the nodule by the vasculature and released into the differentiation zone of the nodule, as is the case for iron (Rodríguez-Haas et al., 2013). There, MtNramp1 introduces iron into the cell (Tejada-Jiménez et al., 2015), while MtZIP6 does the same for zinc, but only in cells that are infected with rhizobia. Once in the cytosol, a fraction of the zinc could cross the symbiosome membrane to supply zinc for the bacteroid. This function has been proposed to be carried out by another ZIP transporter, GmZIP1 (Moreau et al., 2002). However, given the available information on the biochemistry and direction of transport of members of this family (Zhao and Eide, 1996; Lin et al., 2010), it would more likely play a role in buffering zinc content in the symbiosome to prevent toxicity. Instead, a member of the CDF/MTP or P_1B_-ATPase family could be potentially carrying out this role, since they can also transport zinc outside the cytosol, and they have been involved in metal upload to different organelles (Ellis et al., 2004; Eren and Argüello, 2004; Hussain et al., 2004; Desbrosses-Fonrouge et al., 2005). In addition, also working in Zone II, a ZIP or a YSL transporter would be introducing zinc in non-infected cells.

**Figure 7.**
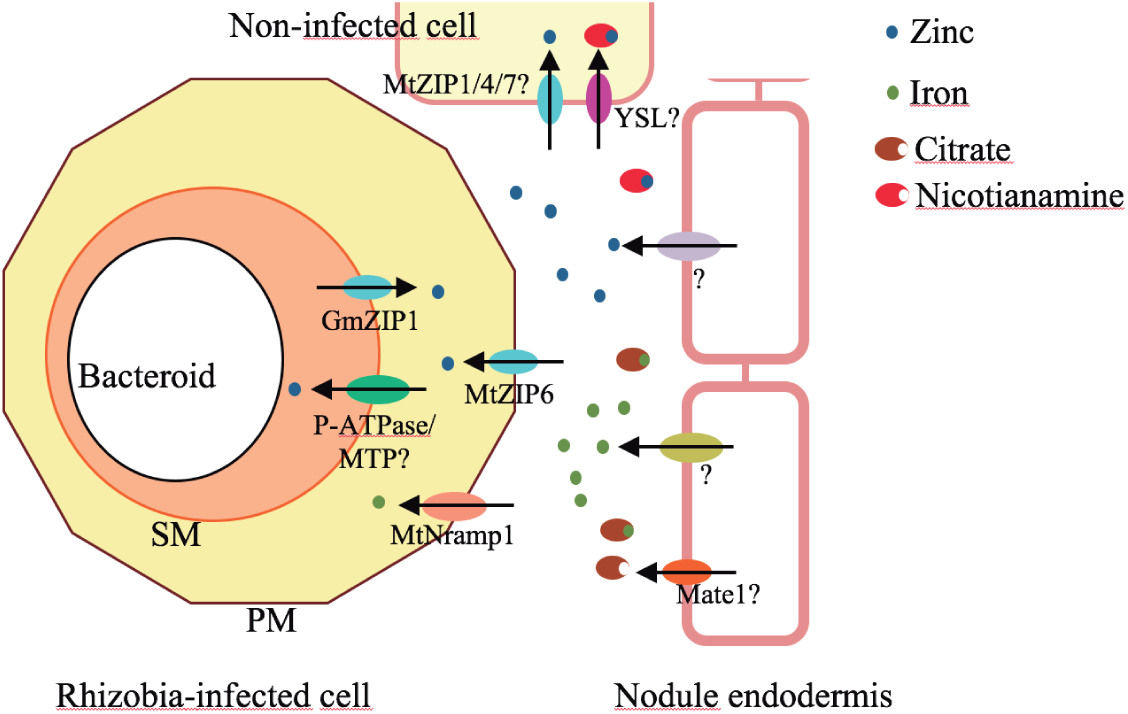
Zinc uptake by nodule cells. Zinc, as iron, is delivered by the vasculature and released in Zone II by yet-to-be-determined transporters. There, MtZIP6 would introduce zinc into rhizobia infected cells, while MtNramp would be responsible for iron uptake. Once in the cytosol MTP, or P-type ATPases would be transporting zinc across the symbiosome membrane. To prevent symbiosome zinc overload, GmZIP1 would be pumping out excess zinc. PM stands for plasma membrane, and SM for symbiosome membrane.

## MATERIALS AND METHODS

### Biological materials and growth conditions

*Medicago truncatula* R108 seed scarification was performed by incubation for 7 min with concentrated H_2_SO_4_. Once washed, seeds were surface sterilized with 50% bleach for 90 s and left overnight in sterile water. After 48 h at 4ºC, seeds were germinated in water-agar plates at 22ºC for 48 h. Then, seedlings were transplanted to sterilized perlite pots and inoculated with *S. meliloti* 2011 or *S. meliloti* 2011 transformed with pHC60 (Cheng and Walker, 1998), as indicated. Plants were cultivated in a greenhouse in 16 h of light and 22ºC conditions, and watered every two days with Jenner’s solution or water, alternatively (Brito et al., 1994). Nodules were collected 28 dpi. Non-nodulated plants were grown in similar conditions of light and temperature but instead of being inoculated with *S. meliloti,* they were watered every two weeks with solutions supplemented with 2 mM NH_4_NO_3_. For hairy-root transformations, *M. truncatula* seedlings were transformed with *A. rhizogenes* ARqua1 carrying the appropriate binary vector as described (Boisson-Dernier et al., 2001).

### RNA extraction and qPCR

RNA was isolated using Tri-Reagent (Life Technologies, Carlsbad, CA), DNase treated and cleaned with RNeasy Minikit (Qiagen, Valencia, CA). Putative DNA contamination on RNA isolation was tested by PCR employing *M. truncatula ubiquitin carboxyl-terminal hydrolase* (*Medtr4g077320.1*) primers (Suppl. Table S1). Denaturing agarose gel was used to verify RNA quality. 1 μg of DNA-free RNA was employed to generate cDNA by using SuperScript III reverse transcriptase (Invitrogen). Gene expression was determined by quantitative Real time RT-PCR (9700, Applied Biosystems, Carlsbad, CA) using primers listed in Suppl. Table S1. The *M. truncatula ubiquitin carboxyl-terminal hydrolase* gene was used to normalize the results. Real time cycler conditions have been previously described (González-Guerrero et al., 2010). mRNA was extracted in three independent experiments, with the threshold cycle (Ct) determined in triplicate. The relative levels of transcription were calculated using the 2^−∆Ct^ method. As control, a non-RT sample was used to detect any possible DNA contamination.

### GUS staining

A transcriptional fusion was constructed by amplifying two kb upstream of *MtZIP6* start codon using primers indicated on Suppl. Table S1, cloned in pDONR207 (Invitrogen) and transferred to pGWB3 (Nakagawa et al., 2007) using Gateway technology (Invitrogen). This led to the fusion of the promoter region of MtZIP6 with the *gus* gene in pGWB3. pGWB3:: *MtZIP6* was transformed in *A. rhizogenes* ARqua1 and used to obtain *M. truncatula* composite root plants as indicated (Boisson-Dernier et al., 2001). GUS activity was determined in 28 dpi plants as described (Vernoud et al., 1999).

### *MtZIP6* knockdown plants

A DNA fragment of the first 469 bp of the *MtZIP6* coding region was amplified by PCR using the primers indicated in Suppl. Table S1, and cloned into the pFRN destination vector (derived from pFGC5941; NCBI accession number AY310901) using Gateway technology (Invitrogen). The DNA fragment is inserted twice in this vector at opposite senses and flanking an intron, so that upon transcription a hairpin double stranded RNA structure is formed. The resulting construct pFRN:: *MtZIP6KD* was transformed into *A. rhizogenes* ARqua1 (Quandt et al., 1993) and used for *M. truncatula* root transformation. The transgenic roots were obtained after kanamycin selection as described Boisson-Dernier et al. (2001). Specificity and efficiency of silencing was checked using real time RT-PCR using the primers indicated in Suppl. Table S1.

### Confocal microscopy detection of MtZIP6-HA

By using Gateway Technology (Invitrogen), a DNA fragment of the full length *MtZIP6* genomic region and the two kb upstream of its start codon, was cloned in the plasmid pGWB13 (Nakagawa et al., 2007). This plasmid fuses three C-terminal HA tags in-frame. Hairy-root transformation was performed as previously described. Transformed plants were inoculated with *S. meliloti* 2011 containing the pHC60 plasmid that constitutively expresses GFP. Nodules collected from 28-dpi plants were fixed by overnight incubation in 4% paraformaldehyde, 2.5% sucrose in PBS at 4ºC. After washing in PBS, nodules were cut in 100 μm sections with a Vibratome 1000 plus (Vibratome, St. Louis, MO). Sections were dehydrated in a methanol series (30, 50, 70, 100% in PBS) for 5 min and then rehydrated. Cell walls were treated with 4% cellulose in PBS for 1 h at room temperature and with 0.1% Tween 20 in PBS for an additional 15 min. Sections were blocked with 5 % bovine serum albumin (BSA) in PBS before their incubation with an anti-HA mouse monoclonal antibody (Sigma, St. Louis, MO) for 2 hours at room temperature. After washing, an Alexa 594-conjugated anti-mouse rabbit monoclonal antibody (Sigma) was added to the sections for 1 h at room temperature. DNA was stained with DAPI after washing. Images were acquired with a confocal laser-scanning microscope (Leica SP8, Wetzlar, Germany).

### Acetylene reduction assay

Nitrogenase activity was measured by the acetylene reduction assay (Hardy et al., 1968). Nitrogen fixation was assayed in RNAi and control plants 28 dpi in 30 ml tubes fitted with rubber stoppers. Each tube contained roots from five independently transformed plants. Three ml of air inside were replaced with 3 ml of acetylene. Tubes were incubated at room temperature for 30 min. Gas samples (0.5 ml) were analyzed in a Shimadzu GC-8A gas chromatograph fitted with a Porapak N column. The amount of ethylene produced was determined by measuring the height of the ethylene peak relative to background. Each point consists of three tubes each with five pooled plants. After measurements, nodules were recovered from roots to measure their biomass.

### Metal content determination

Total reflection X-ray fluorescence (TXRF) analysis was used to determine iron and zinc content in 28 dpi mutant nodules isolated and pooled from 3 sets of 10 different plants each. Analyses were carried out at Total Reflection X-Ray Fluorescence laboratory from Interdepartmental Research Service (SIdI), Universidad Autónoma de Madrid (Spain). Inductively coupled plasma mass spectrometry (ICP-MS) was carried out at the Unit of Metal Analysis from the Scientific and Technology Centre, Universidad de Barcelona (Spain).

### Zinc imaging

Zinc distribution was visualized with the fluorophore Zinpyr-1 as previously reported (Sinclair et al., 2007) with some modifications. Briefly, nodules were hand-sectioned, incubated during 15 min in the dark at room temperature with 200 μM Zinpyr-1 freshly prepared in PBS and washed three times with PBS. Nodule sections were immediately observed under a Leica SP8 confocal microscope using excitation at 488 nm and emission filter at 500-550 nm.

Zinpyr-1 fluorescence quantification was analyzed using FIJI software (v.1.47p). For each section different planes in *z* axis were stacked using Maximum intensity projection, and fluorescence intensity was calculated using “Plot profile” tool from two regions of interest, apical zone (AZ) and fixation zone (FZ), being relativized by the area of each region. Several nodules from independently transformed plants were analyzed to discard variations in the grade of silencing. Representative images for the plants analyzed are showed (Suppl. Fig. 8)

### Bioinformatics

To identify *M. truncatula* ZIP family members, BLASTN and BLASTX searches were carried out in the *M. truncatula* Genome Project site (http://www.jcvi.org/medicago/index.php). Sequences from model ZIP genes were obtained from the Transporter Classification Database (http://www.tcdb.org/) (Saier et al.,2014), NCBI((https://phytozome.jgi.doe.gov/pz/portal.html): *Arabidopsis thaliana* AtZIP1 (*At3g12750*), AtZIP2 (*At5g59520*), AtZIP3 (*At2g32270*), AtZIP4 (*At1g10970*), AtZIP5 (*At1g05300*), AtZIP6 (*At2g30080*), AtZIP7 (*At2g04032*), AtZIP8 (*At5g45105*), AtZIP9(*At4g33020*), AtZIP10 (*At1g31260*), AtZIP11 (*At1g55910*), AtZIP12 (*At5g62160*), AtIRT1 (*At4g19690*), AtIRT2 (*At4g19680*), AtIRT3 (*At1g60960*), AtIAR1 (*At1g68100*), AtZPT29 (*At3g20870*), AtPutZnT (*At3g08650*); *Oryza sativa* OsZIP1 (*Os01g74110*), OsZIP2 (*Os03g29850*), OsZIP3 (*Os04g52310*), OsZIP4 (*Os08g10630*), OsZIP5 (*Os05g39560*), OsZIP6 (*Os05g07210*), OsZIP7 (*Os05g10940*), OsZIP8 (*Os07g12890*), OsZIP9 (*Os05g39540*), OsZIP10 (*Os06g37010*), OsZIP11 (*Os05g25194*), OsZIP13 (*Os02g10230*), OsZIP14 (*Os08g36420*), OsZIP16 (*Os08g01030*), OsIRT1 (*Os03g46470*), OsIRT2 (*Os03g46454*); *Zea mays* ZmZIP1 (*GRMZM2G001803*), ZmZIP3 (*GRMZM2G045849*), ZmZIP5 (*GRMZM2G064382*), ZmZIP6 (*GRMZM2G034551*), ZmZIP7 (*GRMZM2G015955*), ZmZIP8 (*GRMZM2G093276*); *Phaseolus vulgaris* PvZIP1 (*Phvul.001G035800*), PvZIP4 (*Phvul.002G184200*), PvZIP5 (*Phvul.005G048900*), PvZIP6 (*Phvul.005G145900*), PvZIP7 (*Phvul.005G146000*), PvZIP8 (*Phvul.005G149800*), PvZIP9 (*Phvul.006G001000*), PvZIP11 (*Phvul.006G003300*), PvZIP12 (*Phvul.006G055800*), PvZIP13 (*Phvul.006G070200*), PvZIP14 (*Phvul.008G079500*), PvZIP15 (*Phvul.008G259200*), PvZIP16 (*Phvul.008G290500*), PvZIP17 (*Phvul.010G059200*), PvZIP18 (*Phvul.011G058500*), PvZIP19 (*Phvul.002G099700*), PvIRT1 (*Phvul.003G262400*), PvIRT2 (*Phvul.003G262500*), PvIRT3 (*Phvul.009G077700*), PvZIPT (*Phavu. Phvul. L002700*). *Glycine max* GmZIP1 (*Glyma.20G063100*); *Pisum sativum* PsRIT1 (AF065444); *Lycopersicum esculentum* LeIRT1 (AF136579) and *Saccharomyces cerevisiae* ScZRT1 (NP_011259).

Protein sequence comparison and unrooted tree visualization were carried out using ClustalW (http://www.ebi.ac.uk/Tools/msa/clustalw2/), MEGA7 software (http://www.megasoftware.net) and FigTree (http://tree.bio.ed.ac.uk/software/figtree/) using the previously indicated sequences and the newly found *M. truncatula* ones. Previously described *M. trunctula* ZIP1-7 annotation was maintained: *MtZIP1* (*Medtr2g064310*), *MtZIP2* (*Medtr2g097580*), *MtZIP3* (*Medtr3g081580*), *MtZIP4* (*Medtr3g082050*), *MtZIP5* (*Medtr1g016120*), *MtZIP6* (*Medtr4g083570*), *MtZIP7* (*Medtr3g058630*); whereas new MtZIP members were annotated sequentially: *MtZIP8* (*Medtr2g098150*), *MtZIP9* (*Medtr3g081640*), *MtZIP10* (*Medtr3g081690*), *MtZIP11* (*Medtr3g104400*), *MtZIP12* (*Medtr4g065640*), *MtZIP13* (*Medtr5g071990*), *MtZIP14* (*Medtr6g007687*), *MtZIP15* (*Medtr7g074060*), and *MtZIP16* (*Medtr8g105030*).

### Statistical Tests

Data were analyzed by Student’s unpaired *t* test to calculate statistical significance of observed differences. Test results with *P* values less than 0.05 were considered as statistically significant.

## ACKOWLEDGEMENTS

We thank Rosabel Prieto (Universidad Politécnica de Madrid) for her excellent technical support. We would also like to thank Dr. Florian Frugier (Institute of Plant Sciences Paris-Saclay) for kindly sending us the pFRN vector and Dr. Luis M. Rubio (Universidad Politécnica de Madrid) for providing us with an anti-nifH antibody. We acknowledge the Paul Scherrer Institute, Villigen, Switzerland for providing synchrotron radiation beamtime at the microXAS beamline of the Swiss Light Source (SLS).

### List of author contributions

Phylogenetic tree was produced by I. A.; expression studies were carried out by A. S, I. A., V.E, R.C.-R, and M.T.-J.; A.S., R.C.-R., and B.R.-H. determined the tissular and subcellular localization of MtZIP6, phenotypical analyses of RNAi lines were performed by A.S., I.A., M.S., and V.E.; zinc distribution studies were conducted by I.A, C.L., and D.G.; A.S., I.A., and M.G.-G. were responsible for data analyses; J.I., and M.G.-G. were responsible for experimental design, and wrote the manuscript with contributions from all the authors.

### Funding information

This research was funded by a grant from the Spanish Ministry of Economy and Competitiveness (AGL-2012-32974), and a European Research Council Starting Grant (ERC-2013-StG-335284), both to MG-G. RC-R was supported by a FPI fellowship from the Spanish Ministry of Economy and Competitiveness (BES-2013-062674).

